# Live Cell Multicolour Lifetime Imaging Using Genetically Encodable Fluorophores

**DOI:** 10.1101/2022.10.06.511114

**Authors:** Tobias Starling, Irene Carlon-Andres, Maro Iliopoulou, David J. Williamson, Sergi Padilla-Parra

## Abstract

Nine fluorescent proteins (FPs) simultaneous imaging has been demonstrated in a single acquisition using fluorescence lifetime imaging microscopy (FLIM) combined with pulsed interleaved excitation (PIE) for three laser lines. We also show how to unmix spectrally similar FPs in a pixel-by-pixel manner with an analytical non-fitting solution.

## Introduction

Genetically encodable fluorophores, such as the green fluorescent protein (GFP) and its derivatives enabled revolutionary improvements in light microscopy^1^. Over the past 25 years massive protein engineering has been extensively employed to enhance and modify the properties of many fluorescent proteins (FPs)^2^. Methods such as directed evolution, structure-based mutagenesis or recently structure-based rational design^3^ have allowed to reach a colourful FP palette of great variety in terms of both biochemical and spectral properties. One of the most important aspects of FP engineering has been to achieve bright FPs as this will impact both: signal to noise an expression level. The brighter the FP the lower the concentration and therefore less problems with overexpression. The lifetime of a chromophore is defined as the time that electrons take in excited molecules to return to their ground state. It is a unique spectral property that is characteristic of every fluorophore. It is often referred as the spectral fingerprint of fluorophores^4^. Lifetime imaging (FLIM) is independent of the optical path or the fluorophore concentration. Traditionally, it has been employed to reliably quantify protein-protein interactions through Förster resonance energy transfer (FRET) detection. Even if developed a few decades ago, only recently has gathered substantial attention thanks to the development of fast electronics, artificial intelligence, and the decrease of the deadtime of digital detectors capable of photon counting. In practice, this means fastest acquisitions and a better temporal resolution. In addition to these technological advances new efficient and simplified ways of analysing FLIM data popularized FLIM among non experts^5^. When considering FP engineering only a few labs have employed the lifetime as a reference together with brightness, with a few exceptions^6^. Therefore, when consulting the literature regarding lifetime values for existent FPs expressed in live cells little data is available. There is currently no comprehensive list of FP lifetimes under different experimental conditions, especially when expressed in live cells at 37 °C. Here, we have focussed on the use of FLIM to characterize the most appropriate combination of existing FPs for live cell multicolour imaging. We also show that very similar FPs in terms of spectral emission can be separated based on their lifetime information. Current multicolour FLIM approaches are based on combining spectral imaging with lifetime measurements^10,11^. This implies that both the technology and data analysis are too niche/complex to be employed broadly by the biological community. Here, we combine the use of many FPs fully available in public repositories with commercially available technology and lifetime data to demonstrate that the right combination of FPs lifetime, spectral properties and multichannel detection is sufficient to achieve one-shot 9 colour imaging. Importantly this approach is applied to live cells.

## Results and Discussion

The principle of time resolved confocal laser microscopy for lifetime imaging (FLIM) is described in **Figure 1a**. A broad palette of FPs with different spectral properties are excited with three alternatingly pulsed lasers, also termed pulsed interleaved excitation^12^. The lasers were tuned at 440 nm, 485 nm and 594 nm respectively. From a panel of 30 available FPs (**Supplementary Table 1**) individually expressed in live cells, we identified 21 pairwise to hexawise combinations in three spectral channels suitable for multiplexing up to 9 colours in one single acquisition (**Figure 1a)**. For the specific choice of FPs the relative brightness of each fluorophore also needs to be taken into account. Simultaneous excitation with the same laser power for two fluorophores with similar absorption spectra but big differences in brightness will result in the prevalence of the one with higher brightness for their specific spectral channel (**Figure 1b**) regardless their differences in lifetime. Therefore, we have also evaluated the minimal brightness differences between fluorophores to justify our final choice **(Figure 1b)**. The lifetime of each FP considered was analyzed both when expressed individually in live cells (**Supplementary Figure 1**) and when co-expressed together by pairs for each color channel (**Supplementary Figure 2**).

**Figure 1.**
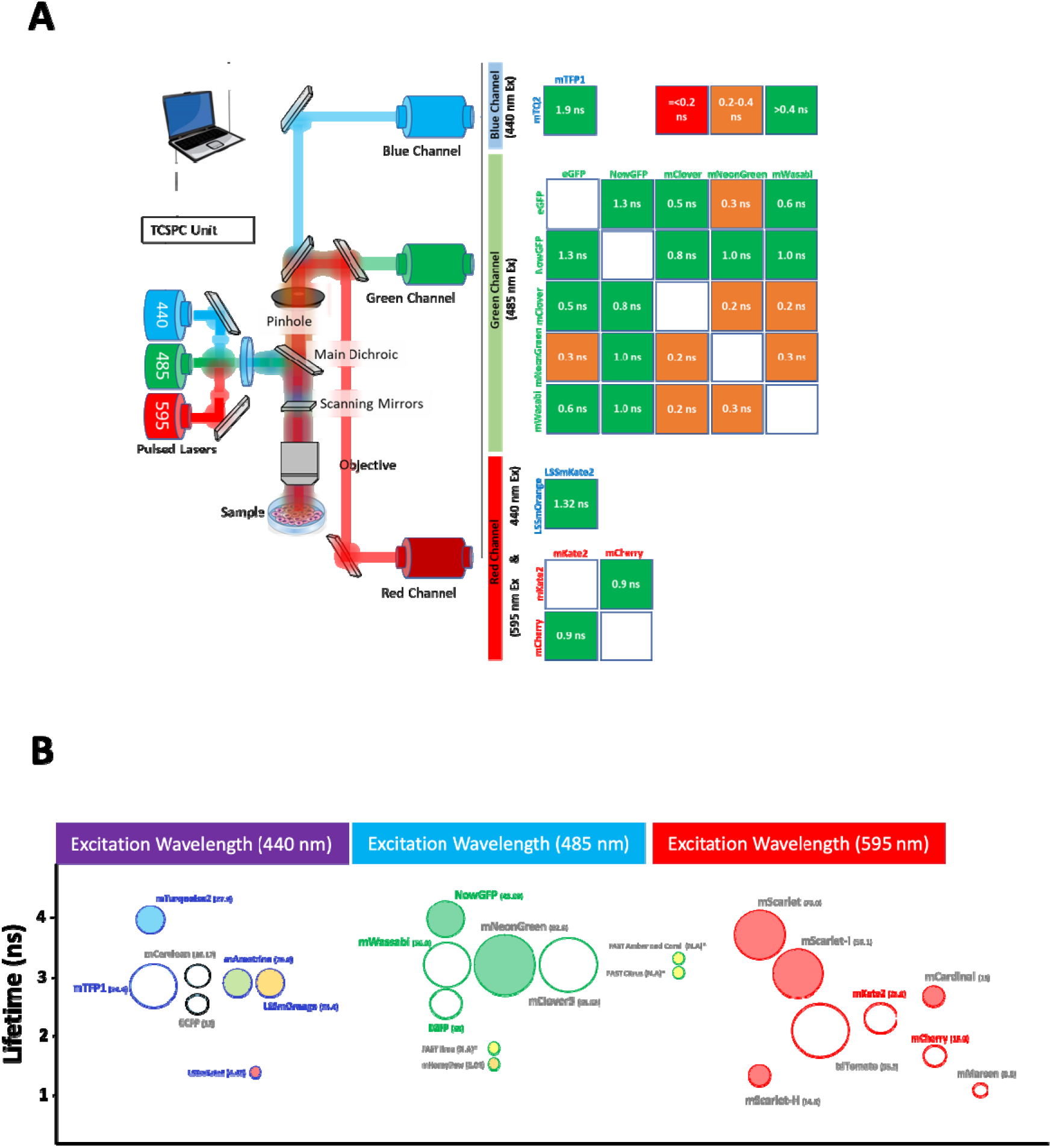
FLIM strategy for multi-color imaging of fluorescent proteins. A commercial time domain time correlated single photon counting (TCSPC) equipped with three pulsed lasers and three detectors (a) was employed. We used two fluorescent proteins per channel with the right spectral, lifetime and brightness (b) properties to achieve nine multicolor imaging in one shot. The main two criteria employed for each channel was the difference in lifetime for similar emission spectral channels (a) and similar brightness as specified in (b).

To validate our non-fitting analytical unmixing approach, two FPs with similar emission spectra and high relative brightness were chosen as a benchmark example: mTFP1^7^ and mTurquoise2^15^. Both proteins have been previously characterized to behave as single exponentials and therefore their separation within the same pixel should be possible assuming a two species model. Employing a simplified FLIM configuration with 440 nm and 485 nm alternating pulsed lasers and one photon counting detector we showed that one can clearly separate both fluorophores when co-expressed in the cytosol of live cells (**Figure 2**). Furthermore, we employ this configuration to ascertain the proportions of the human immunodeficiency virus (HIV-1) receptors (CD4) and coreceptors (CXCR4) in a cell system that mimics the virological synapse (**Figure 2b**). Cells co-expressing CD4-mTFP1 and CXCR4-mTurquoise2 were exposed to cells expressing HIV-1 envelope glycoprotein HXB2 and Gag-mCherry. When looking at cell-cell contacts where Gag-mCherry recruitment where apparent (**Figure 2b**). In these regions, the proportions of CD4-mTFP1 and coreceptor CXCR4-mTurquoise2 for HIV-1 virological synapse^16^ differ as compared to others where these receptors diffuse freely **(Figure 2b)**. Furthermore, we also demonstrate that one can separate pixel by pixel two similar FPs co-expressed together in live cells with our non-fitting analytical solution (**Equation 6–7**) for the green and red channels (**Supplementary Figure 2**).

**Figure 2.**
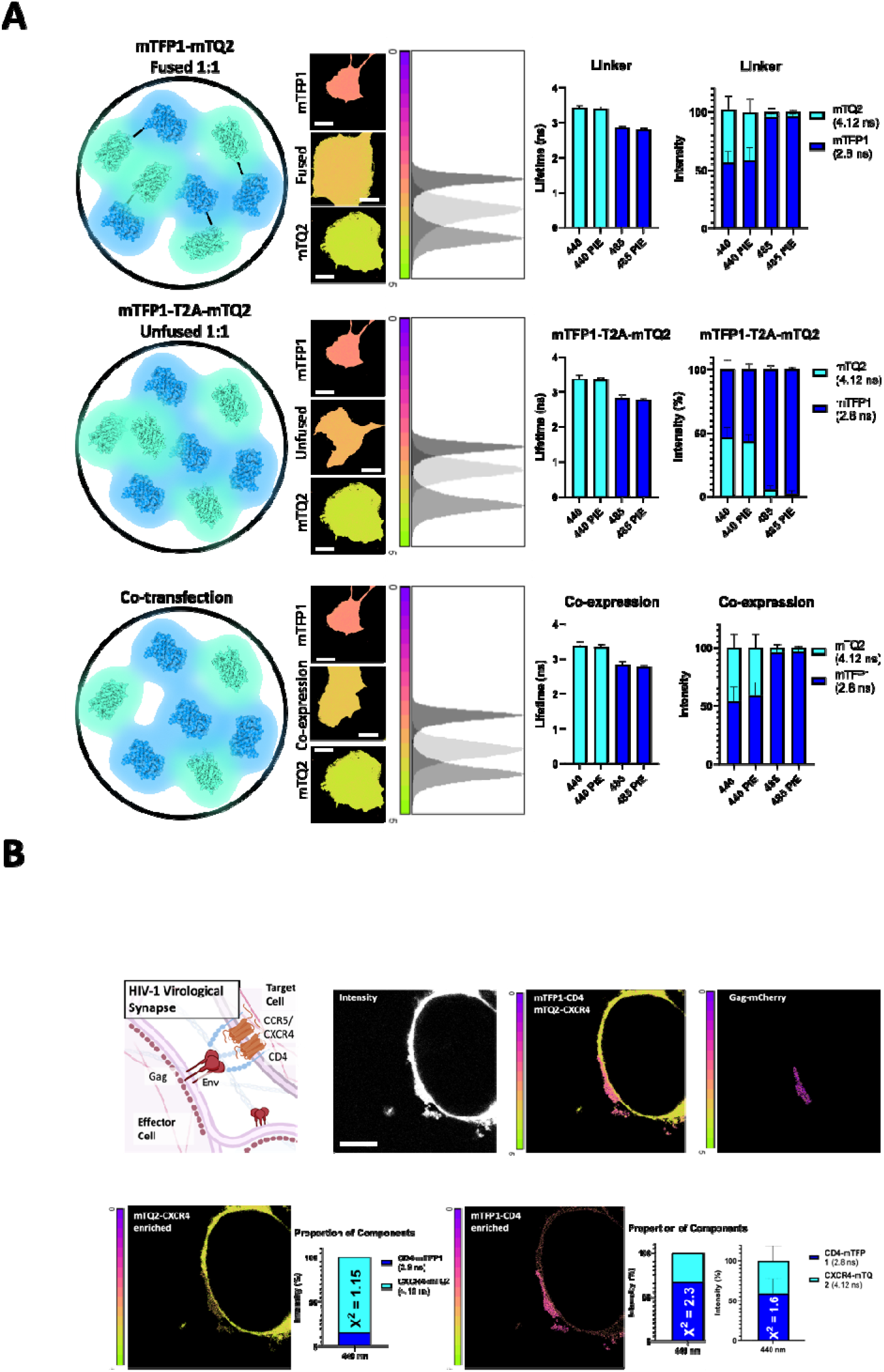
Unmixing of two fluorescent proteins in the same pixel. (a) Live cells (middle panels) expressing a) tandem mTFP1-linker-mTurquoise2 (mTQ2), b) bi-cistronic mTFP1-T2A-mTQ2 and c) co-expression of mTFP1 and mTQ2 were tested to unmix both populations of plasmids. Scale bar 5 μm. Bar charts (right panels) plotting the lifetimes of mTFP1 (dark blue, 2.8 ns) and mTQ2 (cyan, 4.12 ns) when expressed alone were employed as a reference as explained in methods. The percentage obtained for the three cases depicted were 52+/−3%, n = 10; 49+/−4, n = 10 and 50+/−4%, n = 10; respectively. (b) A biological example shows live cells mimicking the HIV-1 virological synapse co-expressing CD4-mTFP1 and CXCR4-mTQ2 (target cells) and HXB2 Env and Gag-mCherry (effector cells). Intensity (left panel) and FLIM images (middle and right micrographs) are depicted showing that one can separate spectrally similar fluorescent proteins. Scale bar 5 μm. Bar diagrams are shown with quantification of the proportion of components colocalizing without Gag-mCherry (bottom left panels) and with Gag-mCherry (bottom right panels). Proportions of CD4-mTFP1 (60+/−10%, n = 5) and coreceptor CXCR4-mTQ2 (40+/−10%, n = 5) differ as compared to other regions where these receptors diffuse freely.

The calculated mean lifetimes^13^ recovered by non-fitting approaches^5^ and therefore recovered instantaneously are utilized to unmix a complex system of up to 9 FPs in one single shot (**Figure 3**). Three different types of HEK293t cells were transfected with 2 to 3 DNA plasmids, then plated together in the same observation well and acquired simultaneously (**Figure 3**). Note that the use of large stokes shift FPs (LSS-FPs) allows to employ the red detector twice (one for the 440 nm line and another for the 594 nm) and increase the number of FPs to be multiplexed. The main caveat of this approach is that most of these LSS-FPs do have low brightness (**Figure 1b**) and therefore their use is limited by the sensitivity of detectors and laser power employed. Despite these limitations we could detect and unmix both Mito-LSS-mKate2 and LSSmOrange^14^ expressed in the cytosol as well as mAmetrine which is also a LSS-FP, but with the maximum emission in the green channel (**Figure 3 and Supplementary Table 1 and 2**).

**Figure 3.**
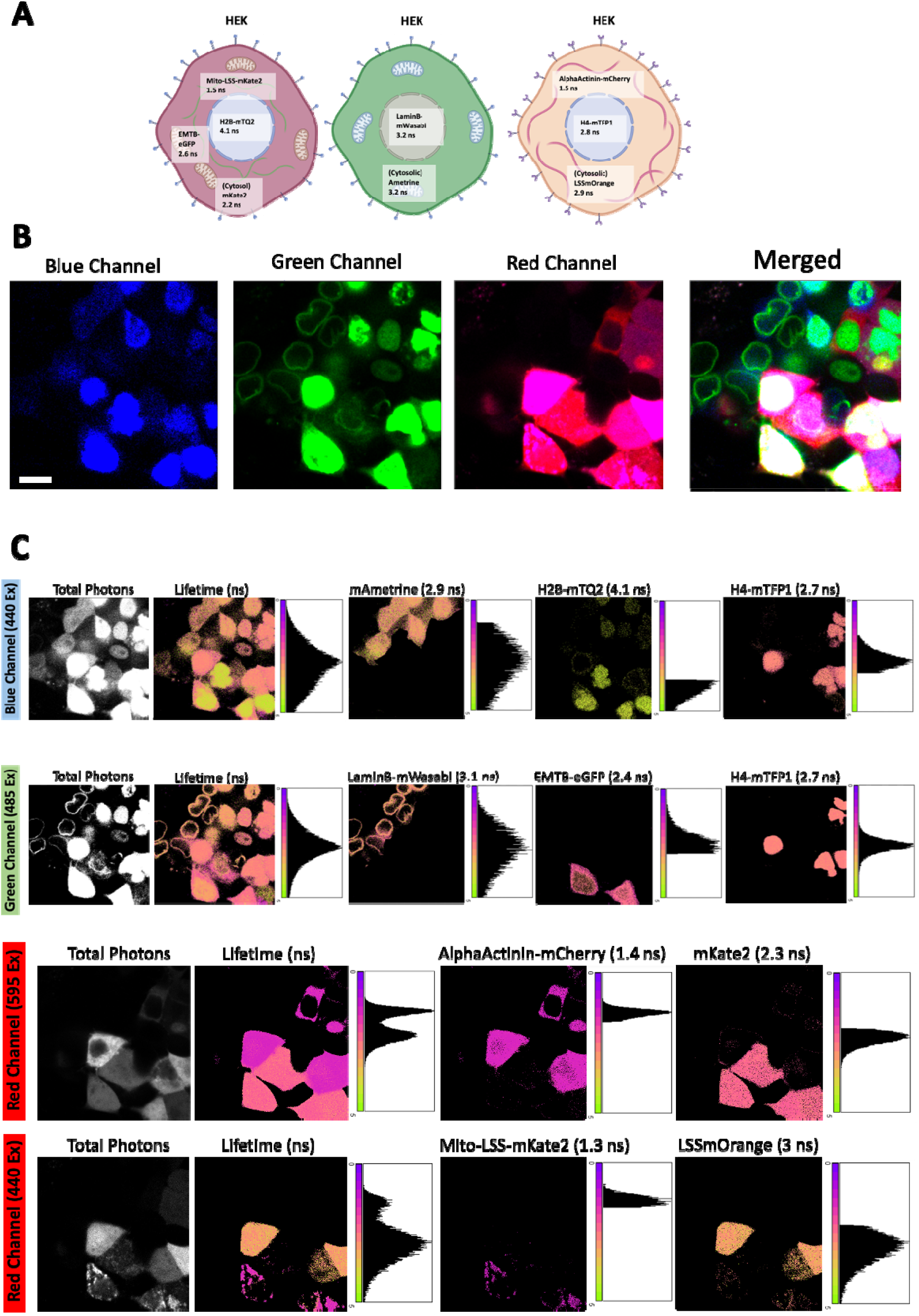
9 color single shot of live cells. Cartoon showing cells co-cultured together co-expressing nine different FP plasmids (three by each cell). The cells were mixed after co-transfection of three different plasmids in each of the three initial wells. (b) The micrographs of live cells expressing nine different plasmids in three different spectral channels are shown. In the blue channel one has cells expressing histone H4 H4-mTFP1 and histone H2B-mTQ2. In the green channel there are cells expressing LaminB-mWasabi, cytosolic mAmetrine and EMTB-EGFP. In the red channel one has cells expressing cytosolic LSSmOrange, Mito-LSSmKate2, Alphactinin-mCherry and Cytosolic mKate2. Scale bar 10 μm. (c) the Intensity micrographs (left columns) together with the overall lifetime images (middle panels) and lifetime unmixed panels (right panels) are depicted. The corresponding pixel lifetime histograms before and after unmixing are also shown. The nine different fluorescent species were separated successfully.

Our multi-color FLIM approach allowed single-shot 9 color imaging of 9 different FPs in live cells (**Figure 3**). This approach permits to image different groups of cells expressing complementary labelled proteins or genetically encodable biosensors. This means that different steps of a given biochemical pathway could be examined and simultaneously imaged with the same conditions in different cells in one shot. This approach also opens the gate to built-in controls imaged at the same time as other dynamic protein interactions. Note that we have only focused on live cell imaging with FPs; but our approach would also be compatible with a combination of FPs and organic dyes and/or biosensors^11^; which would increase drastically the number of colors to be imaged simultaneously. Finally, another pulsed laser could be added, for instance in the far red (635-647 nm) to obtain an extra channel still employing the same red detector. The new developments in far-red FPs would allow to image up two to three more FPs and achieve 12 colors in one single acquistion.

## Methods

### Mammalian cell culture

Untransduced (UT) COS-7 or HEK293t cells and transfected COS-7, HEK293t cells were cultured in Dulbecco’s Minimal Essential Medium (DMEM) (Thermo Fisher, MA) supplemented with 1% penicillin/streptomycin, 10% FBS (Thermo Fisher, MA) at 37°C in a 5% CO2 incubator. Jurkat cells were cultured in RPMI (Gibco, Thermo Fisher, MA) at 37°C in a 5% CO2 incubator.

### Transfection

HEK293 T cells, CV-1 (Simian) in Origin carrying the SV40 (COS-7) UT cells were seeded into an 8 Well Chamber ibiTreat μ-Slide (80826) at 5×10^3 cells/well in 200 μl DMEM and left overnight at 37°C.

Fluorescent proteins were obtained from Addgene (https://www.addgene.org/) transfected using 0.2 ug plasmid DNA. GeneJuice (Sigma-Aldrich, Burlingdon MA, 70967) was used at 3:1 DNA. GeneJuice was mixed with Opti-MEM (ThermoFisher Scientific, MA) at 50 μl/well, vortexed and left at RT for 5 minutes. The GeneJuice+Opti-MEM solution was then added to the 0.2 μg plasmid DNA and left for 15 minutes at RT. This was then added to the 8 well ibiTreat chamber slide and left overnight at 37 °C. Fusion proteins typically required a further day at 37°C before imaging.

### Plasmid constructs

Most of the DNA plasmids used in this article were obtained via Addgene (https://www.addgene.org/; see Supplementary Table 1 and 2). hCXCR4 and hCCR5 were cloned into pmTurquoise2-C1 (#60560) vector by ligating NheI/AgeI fragments into the corresponding sites of the vector, to make hCXCR4-mTQ2 and hCCR5-mTQ2. H2B was subcloned into pmTurquoise2-C1 by ligating HindIII/BamHI fragment into the corresponding sites of the plasmid, making the hH2B-mTQ2 construct. mHoneydew and NowGFP were also cloned into the CD247 (CD3 zeta) plasmid (replacing mEos3.2) from [REF, doi:10.1038/nmeth.3612] Enzymes were purchased from NEB and used according to manufacturers recommendations.

### Image Acquisition

Fluorescence lifetime imaging (FLIM) was performed using the MicroTime200 (Picoquant, Germany, Berlin) time resolved fluorescence microscope. The incubator (Digital Pixel, UK) was preheated to 37 °C for live cell imaging. The sample was excited using pulsed 440 nm, 485 nm or 595 nm diode laser (LDH series Picoquant) with a repetition rate of 20 MHz. The laser beam was coupled to an Olympus IX73 inverted microscope (Japan, Tokio) and focused onto the sample by a 63 ×, 1.2 NA water immersion objective lens (Olympus UPlanXApo). Emission for all figures used a LP488 (Chroma, United Kingdom, London) and detected on a PMA hybrid detector or SPAD (Picoquant). TCSPC was performed using the multiharp 150 (Picoquant). For imaging, the size was 512×512 with a dwell time of 1.3 μs and pixel size of ~0.150 μm/px for 50 frames.

### Average lifetime and unmixing 2 species in the same pixel: a novel non-fitting approach

To unmix with non-fitting approaches and pixel by pixel the fraction of two different FPs an analytical formula was derived which assumes that the decays of the two FPs under consideration could modelled with a single exponential decay.

The kinetics of the fluorescence decay depends on the relative proportion of the different pathways of relaxation once the fluorophores have been excited with the right wavelength. For a single fluorophore in a homogeneous environment the decay can be described as:

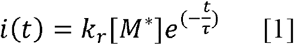

where k_r_ is the constant of relaxation, M the fluorophore, t the time and τ the lifetime. Now, if we consider a two-component system with two different fluorescent proteins then

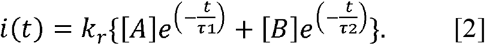

where A is the species with longer lifetime and B the species with shorter lifetime. A discrete double exponential describes the fluorescence decay of both species. If one normalizes the amplitudes of both preexponential factors to 1 we introduce the concept of the fraction each species in a particular pixel:

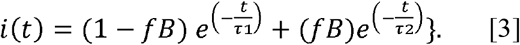

where f_B_ is the fraction of the B component where we assume that the mean average lifetime of B is shorter than the average lifetime of A. Of course, if both lifetimes of the two species are equal one cannot separate their fraction as the overall fluorescence decay will be single exponential (provided that both fluorescent proteins behave as such).

If we introduce the concept of mean lifetime, that with zero background is very similar to the photon arrival time:

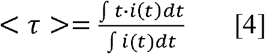

Where t is the time and i(t) the intensity as a function of arrival time.

It can be shown that when considering the above equation as data coming from a discrete sampling one can find the next analytical solution, always assuming a two-component system with two lifetimes:

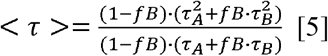

where τ_A_ is the discrete lifetime of the A species, τ_B_ is the discrete lifetime of the B species and τ is the average lifetime. Isolating f_B_ which is the fraction of the B species in the two-component system, from the above expression one obtains the analytical expression to be applied pixel-by-pixel to a FLIM image with two species:

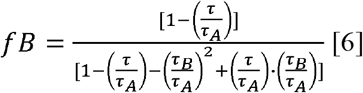

Observe that both τ_A_ and τ_B_ need to be known in advance by performing control experiments. The above expression offers the possibility to obtain the fraction of the species with the shortest lifetime. To obtain the complementary fraction is trivial:

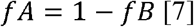

where f_B_ is pre proportion of the B species. All these analytical solutions can be employed pixel-by-pixel.

### Statistics and reproducibility

For FRET-FLIM analyses, calculation was performed with Symphotime software (Picoquant, Berlin) and ImageJ (https://imagej.nih.gov/ij/). Mean and STD for each individual experiment were calculated using GraphPad Prism 9.1.0 software. The sample size and the number of experiment replicate performed are specified in each figure legend.

## Supplementary Figures

**Supplementary Figure 1.**
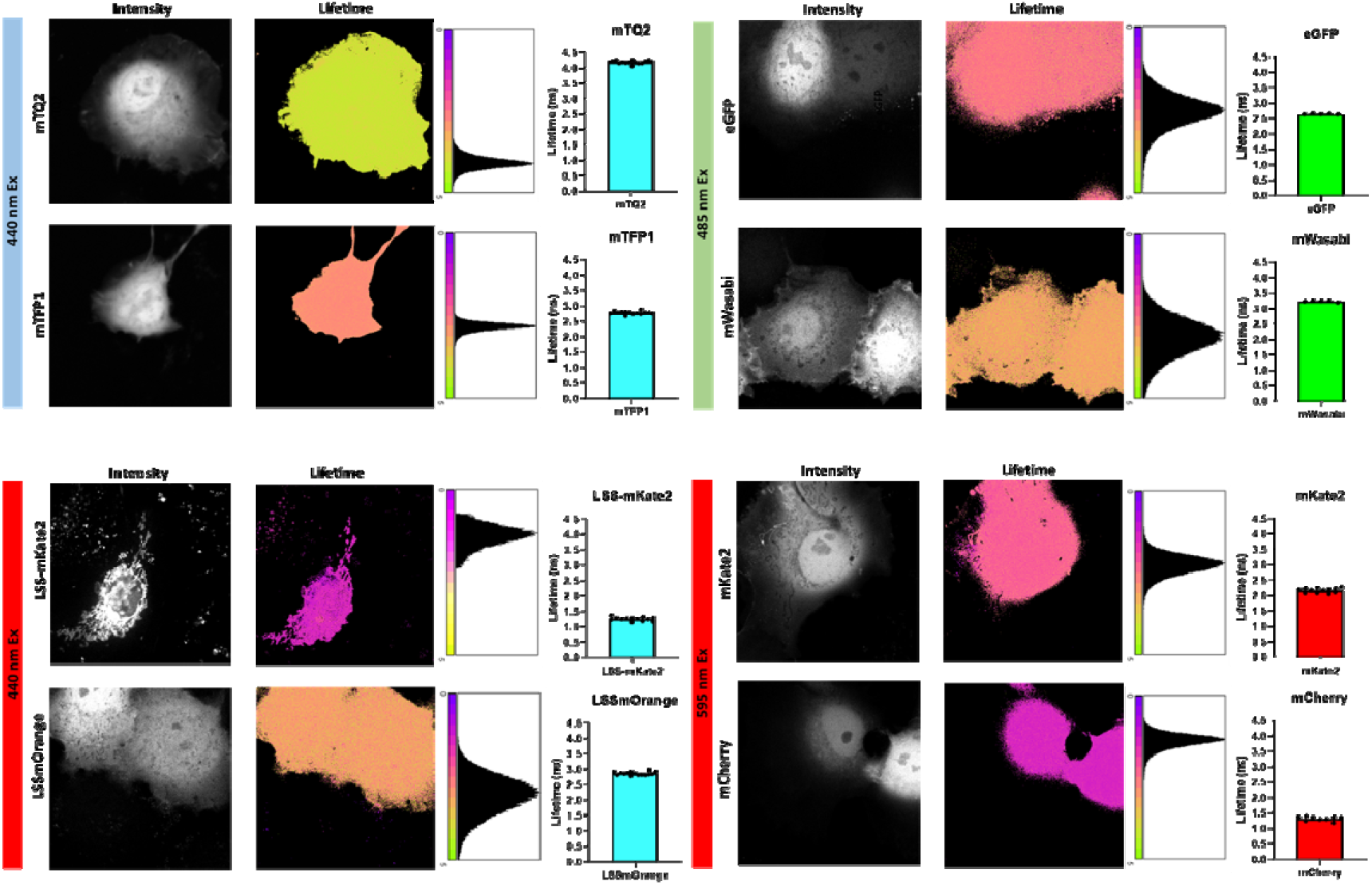
Fluorescent Lifetime determination for each fluorescent protein when expressed in live cells at physiological conditions. A sample of eight different live cells expressing one fluorescent protein species are depicted. Intensity micrographs (Left row) are depicted together with lifetime images (right micrograph) pixel lifetime histograms and bar charts for the average lifetime of each specific fluorescent proteins (n = 5). The blue, green and red channels are shown. Scale bar 5 μm.

**Supplementary Figure 2.**
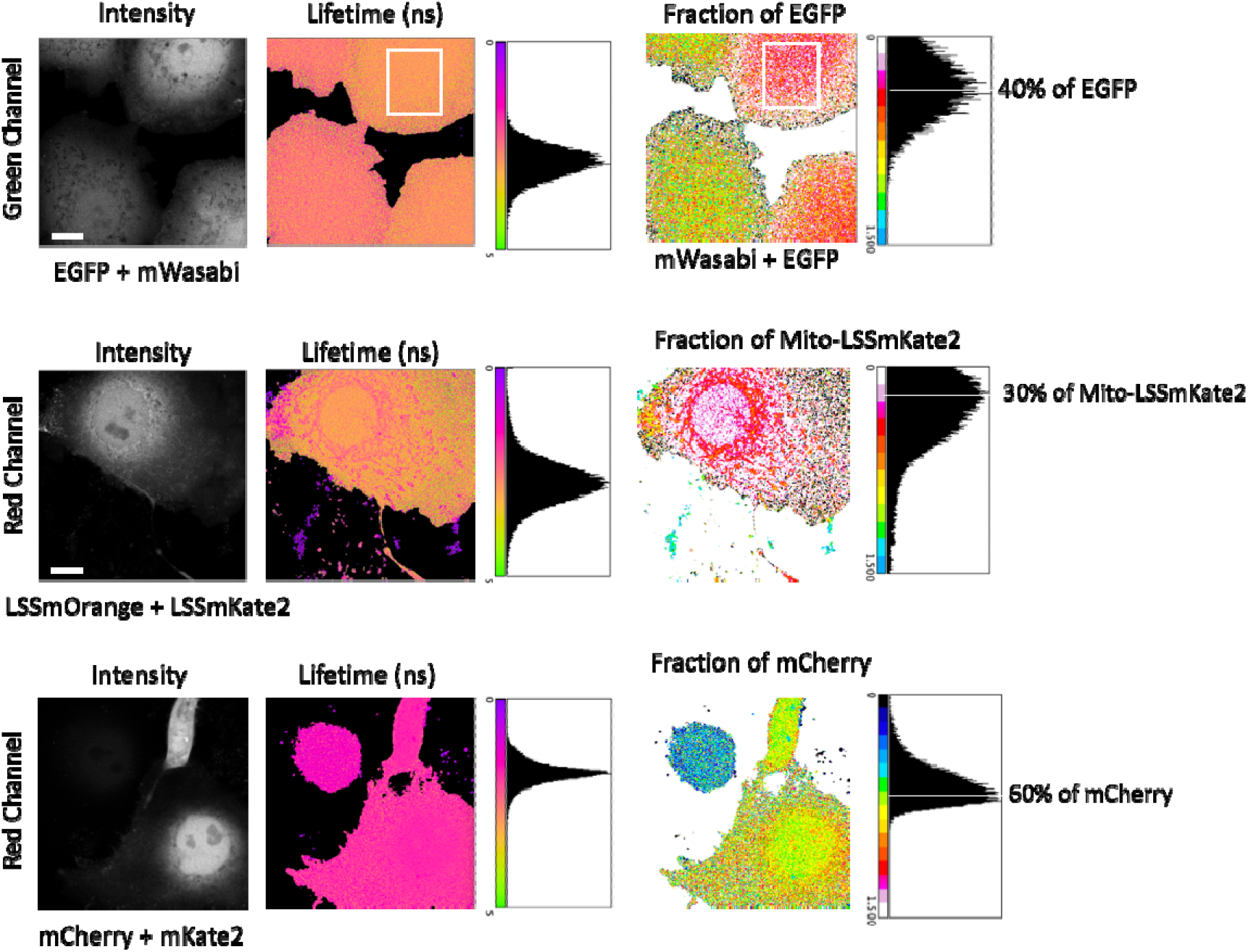
Single pixel pairwise color separation for each channel. Non-fitting analytical solution as described in Methods is demonstrated for green and red channels also (on the top of blue channel, main Figure 2). Intensity images of cells co-expressing either EGFP and mWasabi (green channel), LSSmOrange and LSS-mKate2 (red channel) and mCherry and mKate2 (red channel) are shown. Lifetime images (middle micrographs), pixel histograms and the calculated fraction of fluorescent protein pixel-by-pixel (right micrographs) are also shown in the second and third rows. Scale bar 2 μm. The corresponding histograms are also shown for the average lifetimes and the fraction of the shortest lifetime component. Note that for the fraction of EGFP there were two cells co-expressing EGFP and mWassabi (yellow pixels) and two only expressing EGFP. The application of the analytical formula described in methods clearly separates both. The white region of interest was selected to plot the histogram corresponding to the two-species mixture in live cells.

**Supplementary Table 1.**
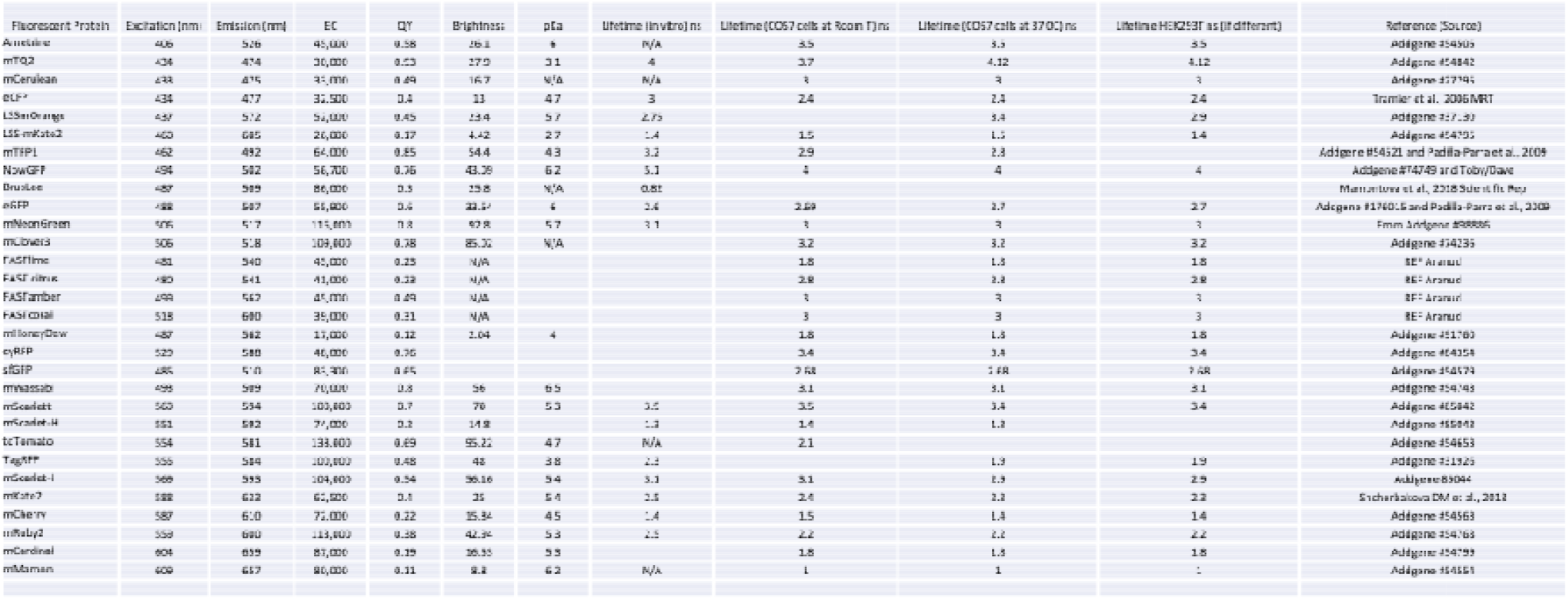
Spectral properties for the list of fluorescent proteins tested when expressed in live cells.

**Supplementary Table 2.**
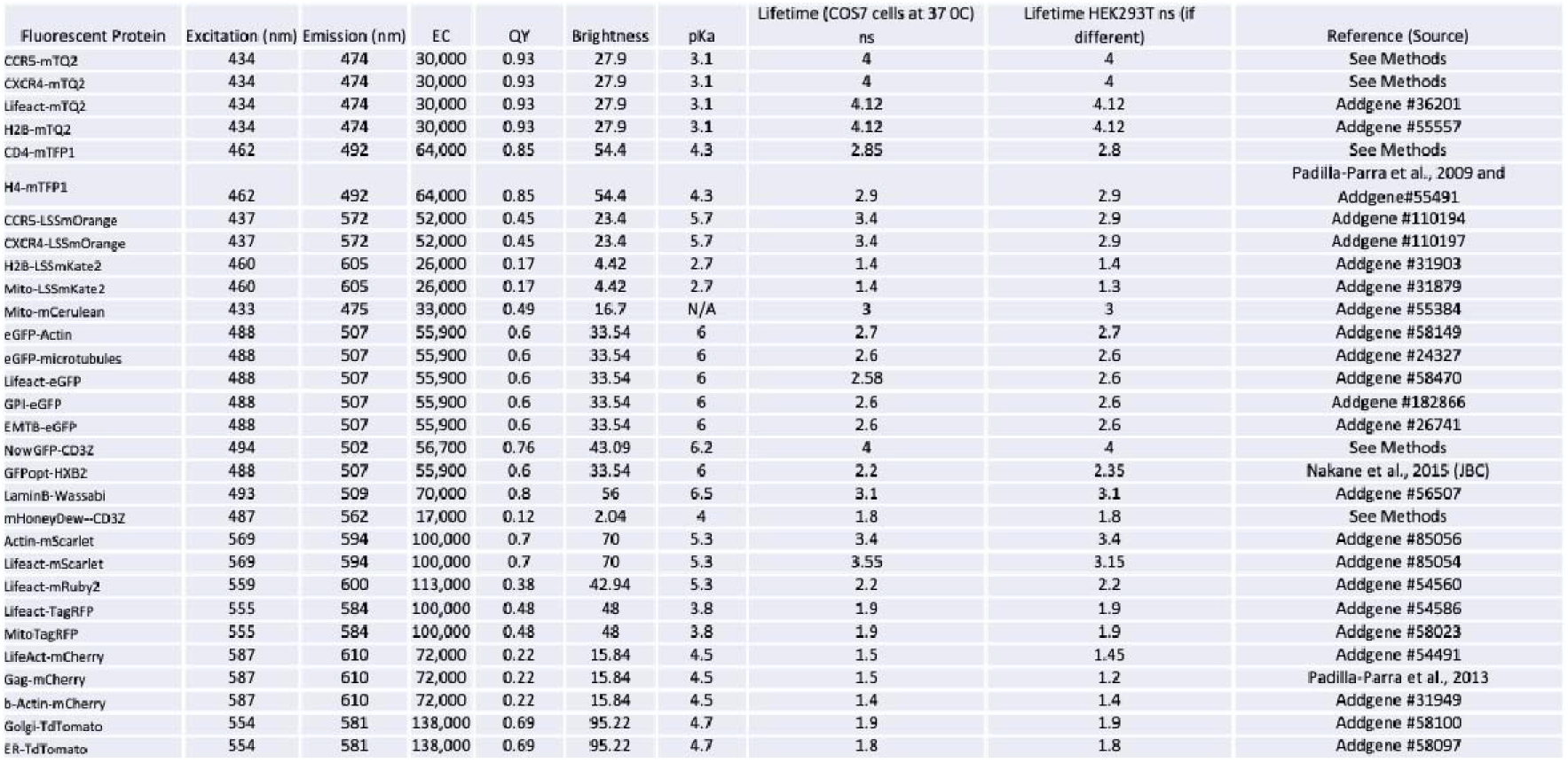
Spectral properties for the list of constructs used labeled with fluorescent proteins tested previously.

## Data availability

(To be updated to a public repository)

## Acknowledgements

We thank all members of the Padilla-Parra lab for valuable discussions and criticism of the paper. This work has been supported by the European Research Council (ERC-2019-CoG-863869 FUSION to S.P.-P.) and the Wellcome Trust Core Award (203141).

## Author Contributions

Conceptualization, S.P.-P.; methodology, T.S, I.C.-A., D.J.W. and S.P.-P.; validation, T.S formal analysis, T.S, D.J.W and S.P-P; resources, S.P.-P.; data curation, D.J.W and S.P.-P.; writing-original draft preparation, S.P.-P.; writing—review and editing, T.S, I.C.-A., M.I, D.J.W. I.C.-A., T.M. and S.P.-P.; visualization, T.S, D.J.W. and S.P.-P.; supervision, S.P.-P.; project administration, S.P.-P.; funding acquisition, S.P.-P. All authors have read and agreed to the published version of the paper.

## Competing interests

The authors declare no competing interests

## Notes

### Competing Interest Statement

The authors have declared no competing interest.

## References

1 Giepmans, B. N., Adams, S. R., Ellisman, M. H. & Tsien, R. Y. The fluorescent toolbox for assessing protein location and function. Science 312, 217–224 (2006). https://doi.org:10.1126/science.1124618

2 Rodriguez, E. A. et al. The Growing and Glowing Toolbox of Fluorescent and Photoactive Proteins. Trends Biochem Sci 42, 111–129 (2017). https://doi.org:10.1016/j.tibs.2016.09.010

3 Hartley, S. M. et al. AlphaFold2 and RoseTTAFold predict posttranslational modifications. Chromophore formation in GFP-like proteins. PLoS One 17, e0267560 (2022). https://doi.org:10.1371/journal.pone.0267560

4 Padilla-Parra, S. & Tramier, M. FRET microscopy in the living cell: different approaches, strengths and weaknesses. Bioessays 34, 369–376 (2012). https://doi.org:10.1002/bies.201100086

5 Padilla-Parra, S., Auduge, N., Coppey-Moisan, M. & Tramier, M. Non fitting based FRET-FLIM analysis approaches applied to quantify protein-protein interactions in live cells. Biophys Rev 3, 63–70 (2011). https://doi.org:10.1007/s12551-011-0047-6

6 Mamontova, A. V. et al. Bright GFP with subnanosecond fluorescence lifetime. Sci Rep 8, 13224 (2018). https://doi.org:10.1038/s41598-018-31687-w

7 Padilla-Parra, S. et al. Quantitative comparison of different fluorescent protein couples for fast FRET-FLIM acquisition. Biophys J 97, 2368–2376 (2009). https://doi.org:10.1016/j.bpj.2009.07.044

8 Tramier, M., Zahid, M., Mevel, J. C., Masse, M. J. & Coppey-Moisan, M. Sensitivity of CFP/YFP and GFP/mCherry pairs to donor photobleaching on FRET determination by fluorescence lifetime imaging microscopy in living cells. Microsc Res Tech 69, 933–939 (2006). https://doi.org:10.1002/jemt.20370

9 Martin, N. et al. Virological synapse-mediated spread of human immunodeficiency virus type 1 between T cells is sensitive to entry inhibition. J Virol 84, 3516–3527 (2010). https://doi.org:10.1128/jvi.02651-09

10 Scipioni, L., Rossetta, A., Tedeschi, G. & Gratton, E. Phasor S-FLIM: a new paradigm for fast and robust spectral fluorescence lifetime imaging. Nat Methods 18, 542–550 (2021). https://doi.org:10.1038/s41592-021-01108-4

11 Niehörster, T. et al. Multi-target spectrally resolved fluorescence lifetime imaging microscopy. Nat Methods 13, 257–262 (2016). https://doi.org:10.1038/nmeth.3740

12 Müller, B. K., Zaychikov, E., Bräuchle, C. & Lamb, D. C. Pulsed interleaved excitation. Biophys J 89, 3508–3522 (2005). https://doi.org:10.1529/biophysj.105.064766

13 Padilla-Parra, S., Audugé, N., Coppey-Moisan, M. & Tramier, M. Quantitative FRET analysis by fast acquisition time domain FLIM at high spatial resolution in living cells. Biophys J 95, 2976–2988 (2008). https://doi.org:10.1529/biophysj.108.131276

14 Shcherbakova, D. M., Hink, M. A., Joosen, L., Gadella, T. W. & Verkhusha, V. V. An orange fluorescent protein with a large Stokes shift for single-excitation multicolor FCCS and FRET imaging. J Am Chem Soc 134, 7913–7923 (2012). https://doi.org:10.1021/ja3018972

15 Goedhart, J. et al. Structure-guided evolution of cyan fluorescent proteins towards a quantum yield of 93%. Nat Commun 3, 751 (2012). https://doi.org:10.1038/ncomms1738

16 Sewald, X., Gonzalez, D. G., Haberman, A. M. & Mothes, W. In vivo imaging of virological synapses. Nat Commun 3, 1320 (2012). https://doi.org:10.1038/ncomms2338

17 Jolly, C., Kashefi, K., Hollinshead, M. & Sattentau, Q. J. HIV-1 cell to cell transfer across an Env-induced, actin-dependent synapse. J Exp Med 199, 283–293 (2004). https://doi.org:10.1084/jem.20030648

18 Iliopoulou, M. et al. A dynamic three-step mechanism drives the HIV-1 pre-fusion reaction. Nat Struct Mol Biol 25, 814–822 (2018). https://doi.org:10.1038/s41594-018-0113-x

